# Protein language models-assisted engineering of Uracil-N glycosylase enables programmable T-to-G and T-to-C base editing

**DOI:** 10.1101/2023.10.29.564583

**Authors:** Yan He, Xibin Zhou, Chong Chang, Ge Chen, Weikuan Liu, Geng Li, Xiaoqi Fan, Yunqing Ma, Fajie Yuan, Xing Chang

## Abstract

Current base editors use DNA deamination enzymes, such as cytidine deaminase (CBE) or adenine deaminase (ABE), to facilitate transition nucleotide substitutions. Combining CBE or ABE with glycosylase enzymes, including CGBE and AYBE, can induce limited transversion mutations. Nonetheless, a critical demand remains for base editors capable of generating alternative mutation types, such as T>G corrections, which could address over 17% of monogenic SNVs responsible for human genetic diseases. In this study, we leveraged protein language models to engineer a uracil-N-glycosylase (UNG) variant with altered substrate specificities to thymines (eTDG). Notably, after only two rounds of testing fewer than 50 predicted variants, more than 50% exhibited a 1.5-11-fold enhancement in enzymatic activities, a success rate much greater than random mutagenesis. When eTDG was fused with Cas9 nickase without deaminases, it effectively induced programmable T to G or T to C substitutions and precisely corrected db/db diabetic mutation in mice (up to 55%). Our findings not only establish orthogonal strategies for developing novel base editors, but also demonstrate the capacities of protein language models for optimizing enzymes without extensive task-specific training data or laborious experimental procedures.

## Introduction

According to ClinVar ^1^, about 58% of human genetic diseases are caused by point mutations. By combining nuclease inactive CRISPR/Cas9 protein and nucleotide deaminase (*i.e.*, cytidine deaminase for CBEs and adenine deaminase for ABEs), base editors (*i.e.*, CBEs ^2, 3^ ^4, 5^ and ABEs^6^) can convert regular bases into modified bases (i.e., C>U or A>I) ^7^. Subsequently, replication over the modified bases induces programmable C>T or A>G base substitutions ^2–9^. These base editors have been widely used in research and pre-clinical studies to correct pathogenic mutations ^10–12^, disrupt gene expression ^13–16^, modulate RNA splicing^17–19^, and diversify genomic DNA sequences^3, 5, 20, 21^. Compared to other gene editing methods, base editors offer the advantage of not provoking DNA double-strand breaks and potential genotoxicities, and only require simple designs to achieve high efficiencies.

Current base editors face limitations in their ability to efficiently induce all types of point mutations, especially transversion mutations. To overcome this limitation, some base editors have incorporated DNA glycosylases, such as N-methylpurine DNA glycosylase (MPG) and Uracil DNA N-glycosylase (UNG) ^22–28^, into CBEs or ABEs. These DNA glycosylases hydrolyze N-glycosidic bonds of the direct products of ABE or CBE (dI or dU, respectively) to generate abasic sites. If not repaired promptly, DNA polymerase predominantly adds cytidine opposite to an AP site in eukaryotic cells (C rule) in eukaryotic cells^29^, enabling the creation of C>G or A>G mutations. However, the incorporation of additional modules to already bulky base editors increases the difficulties of efficient *in vivo* delivery and the risk of off-target editing by both the deaminase and DNA glycosylases. Furthermore, the editing outcomes of these transversion base editors involve two-step reactions, which make it difficult to predict the final outcome accurately ^25, 30^.

Improving gene editing modules based on natural enzymes often relies on directed evolution and structural-assisted engineering. However, these approaches typically demand substantial task-specific training data and extensive experimental efforts. In contrast, protein language models (PLMs) ^31, 32^ have been developed by learning to predict the masked amino acid in a protein sequence, given the surrounding amino acids. PLMs have been utilized to predict the direction of natural evolution ^33^, facilitate the design of novel proteins ^34^, and improve affinities of human antibodies ^35^. Yet, optimizing enzymes presents additional challenges than other proteins (*e.g.,* proteins) using PLMs. While antibodies primarily rely on the biophysical properties of their complementarity-determining regions (CDRs) for antigen binding, optimizing enzymes requires enhancement of both their biophysical and biochemical properties of the intricate protein structures, many of which are poorly understood. Moreover, enzymes exhibit remarkable diversity in terms of their classes and catalytic mechanisms, and each enzyme may require specific engineering approaches. Therefore, the potential application of PLMs pre-trained with all protein sequences remains uncertain in selecting the most promising enzyme variants.

In this study, we harness Uracil N-glycosylase variants to create orthogonal transversion base editors (CGBE and TSBE) independent of deaminases. We also developed a strategy based on protein language models and engineered an enhanced Uracil N-glycosylase variant with specificities toward thymines (eTDG). By testing a limited number of variants, we obtained eTDG with significantly improved activities (>2-fold increase). Integration of eTDG into spCas9 protein resulted in the development of TS(G or C)BE, a highly efficient tool for inducing programmable T>G or C substitutions, and precisely corrected an obesic mutation (db/db G>T) mouse embryos. This work established novel base editors and showcases the potential of utilizing protein language models to streamline the engineering of gene editing tools with enhanced specificities and minimal experimental burden.

## Results

### Fusion of a UNG variant targeting cytidine with nCas9 enables programmable C>G base editing

Given the limitations of previously reported transversion base editors (**Suppl. Figure 1A**), we reasoned that an alternative approach is to directly target normal bases on DNA using an engineered glycosylase with altered substrate specificities, which could potentially enable the one-step creation of abasic sites and the design of more efficient and specific transversion base editors (**Figure 1A**).

**Figure 1.**
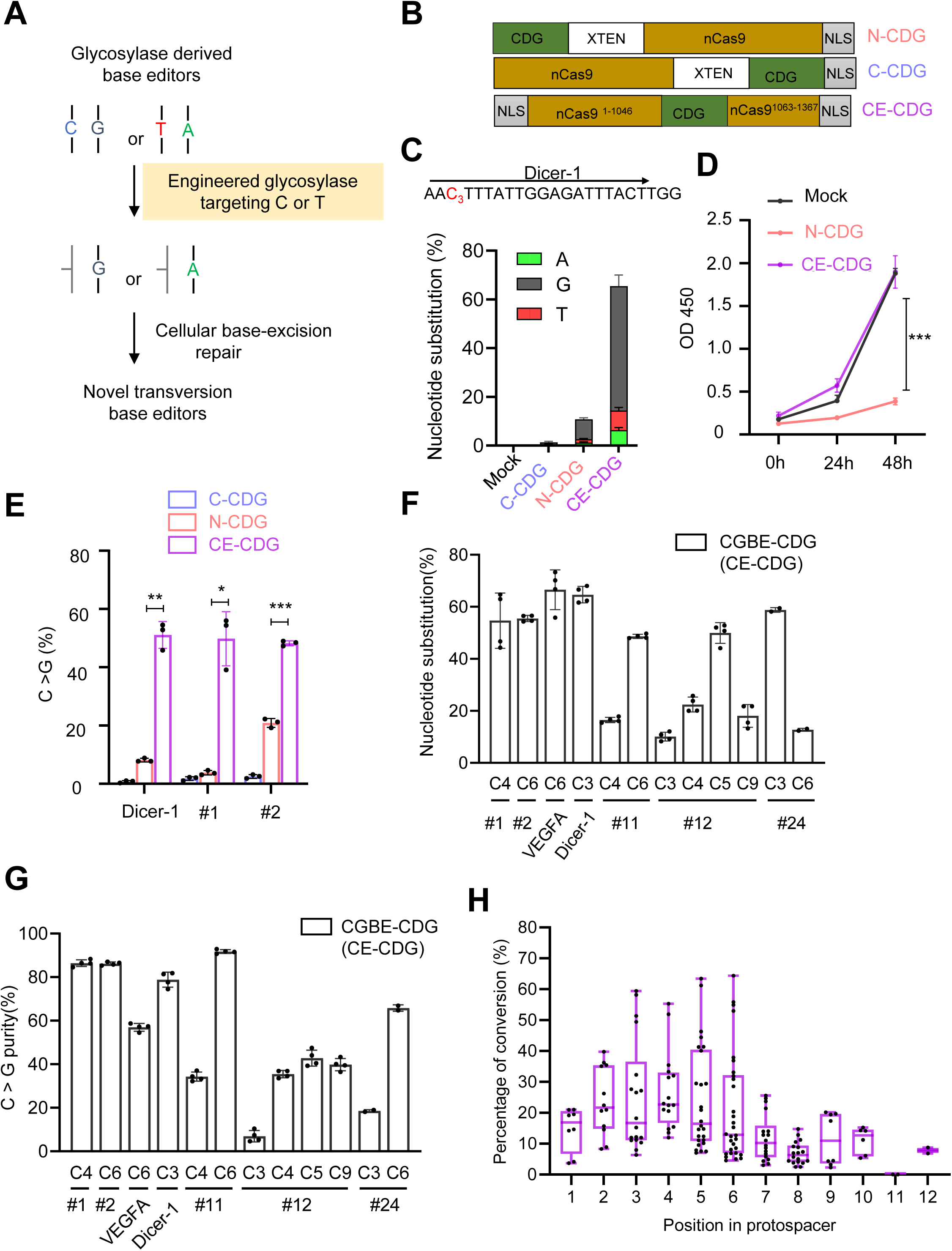
Embedding a variant of UNG2 within Cas9 nickase enables programmable C>G transversion base editing. (**A**) Schematic of inducing transversion mutations using engineered DNA glycosylases targeting normal nucleotides. Glycosylase variants with altered substrate specificities (e.g., UNG variants targeting cytidine or thymine) can be fused with Cas9 nickase protein (nCas9 D10A) to enable one-step generation of abasic sites and potential transversion mutations without DNA deamination. (**B**) Diagrams showing CGBEs based on a variant of hUNG (hUNG N204D) targeting cytidines. NLS, nuclear-location sequences; Cas9n, SpCas9 nickase (D10A); CDG, UNG variant (hUNG N204D) with cytidines as substrates. CE-CDG, Cas9-embedded CDG; N-CDG, CDG was fused to the N-terminus of nCas9 protein; C-CDG, CDG was fused to the C-terminus of nCas9 protein. (**C-E**) Embedding CDG within the nCas9 protein improved editing efficiencies (**C**) and suppressed cytotoxicity (**D**). HeLa cells were transfected with nCas9-CDG fusion proteins with indicated configuration as in (**B**) and a sgRNA targeting the Dicer-1 locus. (**C**) Five days after the transfection, an amplicon covering the protospacer was amplified from the genomic DNA and was analyzed by high-throughput sequencing. Nucleotides with detectable mutations (mutant frequency > 0.1%) were depicted. The percentages of reads containing C>G, A, or T conversion were indicated. Locations of the sgRNA and the PAM sequence are shown on the top of the target sequence. Data are representative of three independent experiments. Error bars stand for the standard variation of the mean. (**D**) The proliferation of HeLa cells transfected with indicated constructs was determined by CCK8 assays. Data are representative of three independent experiments. Error bars stand for the standard deviation of the mean. ***, p<0.001 in Student’s *t* tests. (**E**) as in (**D**), efficiencies of inducing C>G substitution using indicated nCas9-CDG configuration together with Dicer-1, #1, or #2 sgRNAs. Data are summary of three independent experiments. (**F-G**) Summary of activities of CE-CDG in seven endogenous loci. As in (**E**), base substitution (**F**) and C>G conversion purity (**G**) in seven endogenous loci were determined by HTS. Note the mutated nucleotides were labeled as their position relative to the 5’ end of the protospacer. No mutations were observed beyond the protospacers. (**H**) The editing window for CE-CDG was determined by plotting the frequencies of cytosine editing across various protospacer positions, with the nucleotide furthest from the PAM designated as position 1. Box plots represent the interquartile range (25th to 75th percentile), horizontal lines indicate the median (50th percentile), and whiskers extend to the minimum and maximum values.

Prior investigations into uracil DNA glycosylase (UDG2) have shown that two human UDG2 variants (N204D or Y147A) can switch their substrate specificity from uracil to cytosine or thymine, denoted as cytidine DNA glycosylase (CDG) or thymine DNA glycosylase (TDG), respectively ^36–38^ ^39^. Initially, we attempted to fuse CDG to either the N- or C-terminus of nickase SpCas9 (nCas9) protein (**Figure 1B**); however, this resulted in only minor C>G substitutions (less than 10%) at the Dicer-1 locus when transfected into HeLa cells together with a sgRNA (**Figure 1C**). Notably, transfection of these editing modules led to cytotoxicity as evidenced by reduced cellular proliferation (**Figure 1D**).

To minimize the cytotoxicity, we adopted an embedding strategy by inserting CDG into the middle of the nCas9 protein between F1046 and I1063 ^40^. While embedding the glycosylase in nCas9 may hinder its free access to off-target sites and reduce its toxicities, it does not prevent access to on-target sites due to forced tethering via nCas9 and sgRNA^40, 41^. Indeed, embedding CDG within the nCas9 protein (named CGBE-CDG) decreased cytotoxicity to levels similar to vector transfection (**Figure 1D**). This strategy also increased the editing efficiencies ac,hieving over 50% C>G substitutions at protospacer position 3 of Dithe cer-1 site (counting the protospacer adjacent motif [PAM] as positions 21–23) (**Figure 1C**). Similar enhancement was observed in the other two endogenous loci (**Figure 1E**). The increased editing efficiencies were likely because the transfected cells with the highest base editor activities did not die from sgRNA-independent activities of the glycosylases against cytidine. Consequently, frequencies of indels, which were likely caused by CDG activities beyond the editing window, were decreased in cells transfected with CGBE-CDG than in cells transfected with N-CDG (**Suppl. Fig.1B**). Moreover, the activities of CGBE-CDG were dependent on the CDG enzymatic activities since co-expression of UGI, a phage-derived UNG2 inhibitor that blocks UNG2/DNA interactions ^42^, inhibited the activities of CGBE-CDG in inducing base substitutions (**Suppl. Fig. 1C**).

Further examination of CGBE-CDG activities in seven endogenous loci revealed over 50% (56.8 ± 8.3%) nucleotide substitution with C>G purities exceeding 70% (72.7 ± 18.5%) (**Figure 1F-G, Suppl. Fig. 2**). CGBE-CDG primarily induced base substitutions at cytidines located at positions 2-6 within the protospacers (**Figure 1H**), and recognizes a loose CW(A/T)S(G/C)motif (**Suppl. Fig.1D**). Compared to previously developed CGBEs ^23–26^ (**Suppl. Fig.3A**), CGBE-CDG showed comparable or higher activities (**Suppl. Fig.3B-C**) along with reduced C>T byproducts (**Suppl. Fig.3D**). However, in some loci, CGBE-CDG induced a higher frequency of indels (**Suppl. Fig.1E, Suppl. Fig.3E**). Analysis of off-target editing also revealed nucleotide substitution levels below 0.20% at predicted off-target loci (**Suppl. Table1**). These results collectively show that the CDG and nCas9 fusion protein can effectively induce the programmable C>G base editing with the guidance of sgRNAs. Furthermore, these findings underscore the potential of engineering glycosylases to serve as orthogonal base editors for transversion base editing.

### Fusion of a UNG variant targeting thymine with nCas9 enables moderate T>G or T>C base editing

Next, we aimed to develop glycosylase-based base editors that target thymine, a capability that has not been previously achieved with deaminase-based base editors, while T>G correction could fix over 17% of monogenic SNVs associated with human genetic diseases^43, 44^. Fusion of the TDG (UNG2 Y147A) with nCas9 either internally or at the terminus (**Figure 2A**), however, only led to less than 10% substitution at the T5 position of the Dicer-1 locus (**Figure 2B-C**). In addition to T>G conversion, T>C substitutions were another primary editing outcome of the TDG-nCas9 fusion protein. Thus, we named the TDG internally embedded into nCas9 protein as T to S (G/C) base editor (TSBE1). Optimization of linkers connecting TDG and nCas9 to five residues moderately improved editing efficiencies to around 12% at T5 of Dicer-1(12.7 ± 1.28%) (**Figure 2D-F**, named as TSBE2). Further examination of TSBE2 activities in 12 endogenous loci revealed over 10% (10.7 ± 7.2%) nucleotide substitution with T>S purities exceeding 84% (84.3 ± 8.3%) (**Figure 2G, Suppl. Fig. 4A**). TSBE2 produced higher indels when there are multiple T’s within the editing range (**Suppl. Fig. 4B**). TSBE2 primarily induced base substitutions at cytidines located at positions 3-8 within the protospacers (**Suppl. Fig. 4C**), and TSBE2 didn’t have a preference over nucleotide sequences surrounding the targeted Ts (**Suppl. Fig. 4D**). Similar to CGBE-CDG, activities of TSBE2 were effectively inhibited by co-expressing UGI (**Figure 2H**), confirming that the glycolysis activity of TDG was essential for the editor’s function.

**Figure 2.**
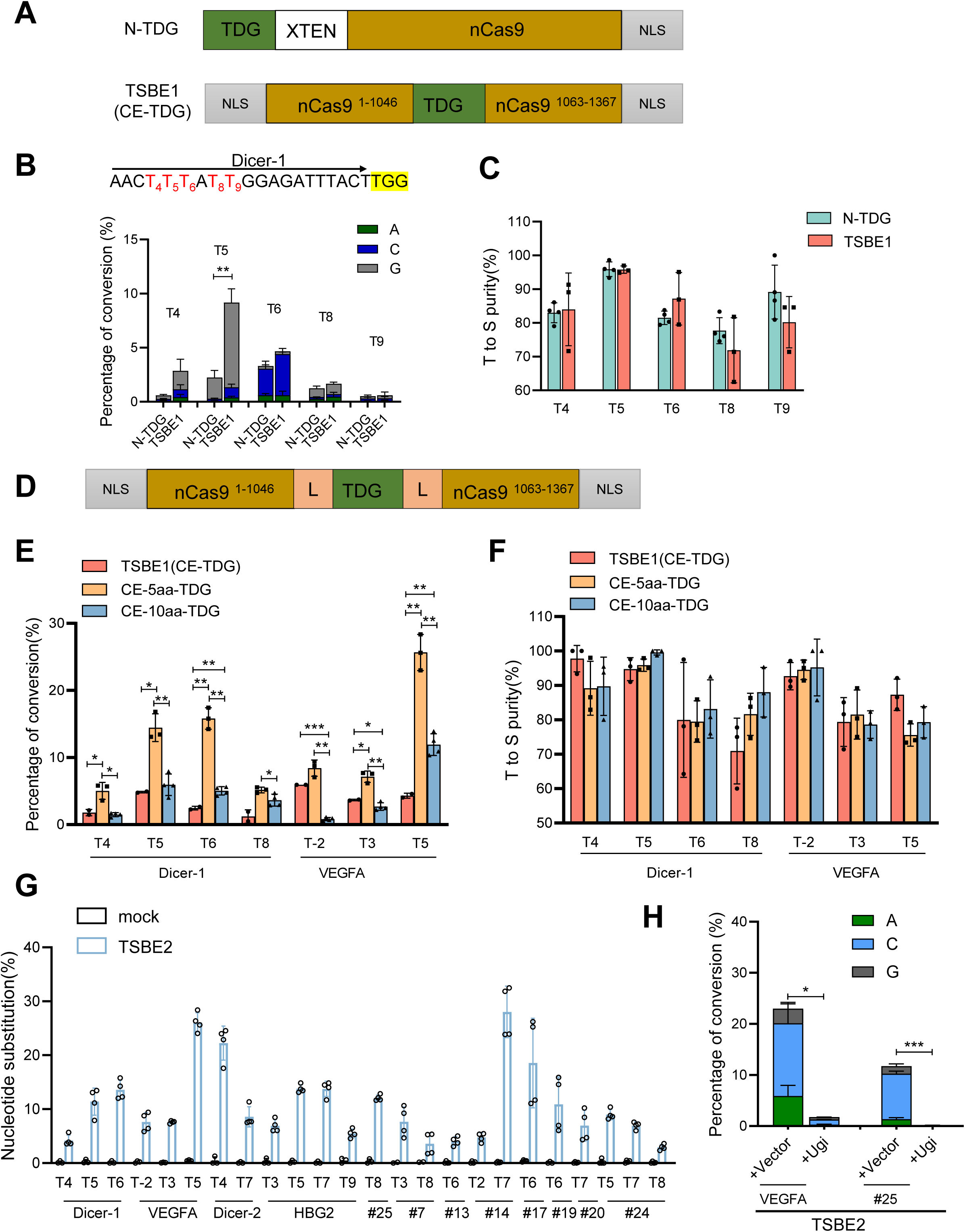
Embedding of TDG in nCas9 generates moderate T>G or T>C conversion. (**A**) The schematic illustrates the configuration of TDG fused to either the N-terminus or middle of the nCas9 protein. TDG refers to the UNG Y147A variant, XTEN represents the XTEN linker sequence, nCas9 denotes the SpCas9 nickase, and NLS stands for the nuclear localization sequence. (**B-C**) Higher nucleotide substitution induced by TSBE1 (CE-TDG) than by the N-terminal fused TDG at the Dicer-1 locus. (**B**) A diagram depicting the sequences of the protospacer and PAM of Dicer-1 sgRNA, with targeted bases highlighted and numbered from the distal end of the protospacer to the PAM sequence. TSBE1 and N-terminal fused TDG were transfected into HeLa cells along with Dicer-1 sgRNA. After seven days, nucleotide substitutions at the indicated thymines were determined through high-throughput sequencing (HTS). The data represented are summary of three independent experiments, and the error bars represent the standard deviation of the mean. (**C**). As in (**B**), HTS analysis of T to S (G+C) purities at the indicated thymines. (**D-F**) Inclusion of a 5 amino acid linker between TDG and nCas9 enhances editing activities. (**D**). A Diagram showing the embedding TDG into nCas9 protein, L, a five or ten amino acid linker. Similar to (**B**), HeLa cells were transfected with TSBE1 (no linker) or TSBE (with a five or ten amino acid-linker) and sgRNAs targeting the VEGFA or Dicer-1 loci. After seven days, nucleotide substitution (**E**) and T>S purity (**F**) were determined via HTS. The data shown are summary of three independent experiments, and the error bars represent the standard deviation of the mean. (**G**) Summary of editing efficiency of TSBE2 in 12 endogenous loci. HeLa cells were transfected with the indicated sgRNA and TSBE2 at a 1:1 molar ratio. Seven days after the transfection, nucleotide substitution was determined with HTS. (**H**) The enzymatic activities of TDG are essential for TSBE to induce nucleotide substitutions. HeLa cells were transfected with TSBE2 and VEGFA or #25 sgRNA, with or without UGI. After seven days, the editing efficiencies were determined using HTS. The data presented are summaries of two or three independent experiments.

### Protein language model facilities design of efficient TDG variants

To further enhance the activities of TSBE2, we employed protein language models (PLMs) ^31, 32^ and adopted the Protein language Model ESM (Evolutionary Scale Modeling), which was trained with multiple datasets, including Uniprot, Uniref50, Uniref100, SwissProt, etc^31, 45–47^. The training objective of ESM is to predict the masked token, which is a word in the natural language process task and an amino acid for the protein sequence ^48–52^. To find high-fitness enzyme variants, all the potential mutation sites within the wild-type sequences were inputted into an ESM, which outputs the corresponding likelihood of the mutation site through a Masked Language Model head - a module designed to transform a representation from the ESM into a desired format. We selected 17 different ESM models and combined them with nine ranking strategies (**Suppl. Table 2**) to perform benchmarking on a previously published dataset ^53^, which includes the folding stabilities of all single residue variants of 542 proteins. We found that the combination of the Wildtype Marginal Probability (esm1v_5) and the esm2_t33_650M_UR50D model performed the best in the precision of predicting the function-enhancing variants, such that more than 45% of the selected mutations had a fitness above the wildtype (**Suppl. Fig. 5A and Suppl. Table 3**).

To engineer a TDG variant with enhanced activities using PLM, we determined the fitness of all single-residue substitutions of TDG (UNG2 Y147A) (**Figure 3A**). As anticipated, the UNG2 A147Y variant, which reverted the substrate-altering mutation, achieved the highest ranking (**Suppl. Table 4**). Among the top 50 variants, 40% was concentrated in the N-terminal tail (residue 1-92) with a density of 0.22 variants per residue, in contrast to the catalytic domain (residue 93-304) with a density of 0.13 variants per residue (**Suppl. Figure.5B-C**). This bias may arise from the fact that the catalytic domains are highly conserved, leaving limited room for enhancing their evolutionary fitness. For experimental validation, we selected the top 38 variants (excluding A147 mutations) for insertion into the nCas9 protein. In the Dicer-1 site, 33 out of the 38 variants exhibited superior activities compared to WT TDG, as determined by the percentage of T>G substitutions at T5 and overall editing efficiencies in the Dicer-1 locus (**Figure 3B**). Sixteen tested variants demonstrated more than a 1.5-fold improvement, and two variants (L74Q and G107E) exhibited over a 2-fold enhancement in editing efficiencies.

**Figure 3.**
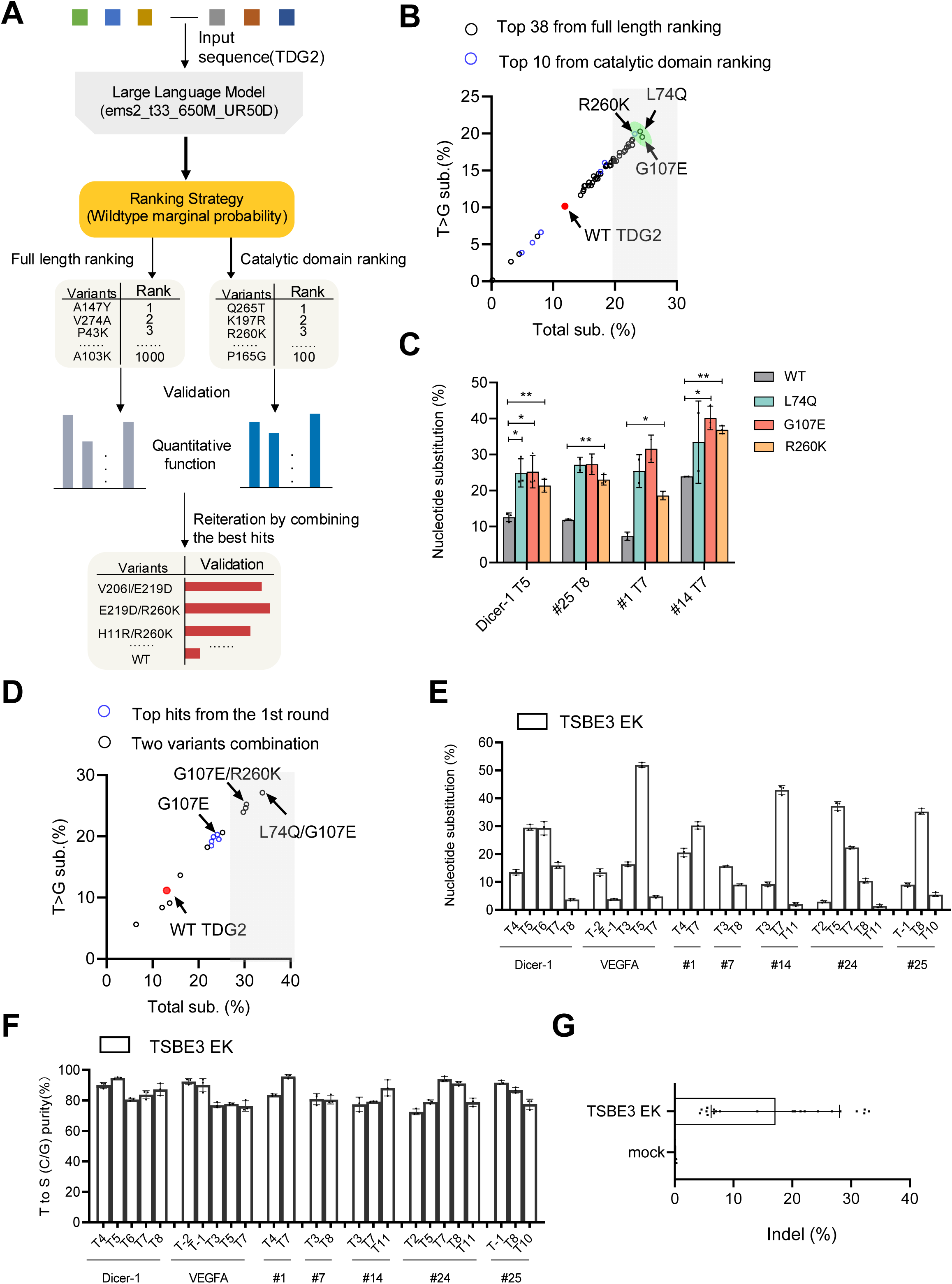
A strategy based on protein language models facilitates generation of efficient TDG variants for inducing T-to-G and T-to-C conversion. (**A**) Schematic of applying protein language models (PLM) to engineer UNG variants with enhanced activity against thymines. The fitness of all single-residue variants of TDG was assessed using the esm2_t33_650M_UR50D model and ranked employing the Wildtype Marginal Probability strategy. The top-ranked 38 variants were selected for experimental validation after being inserted into nCas9. Concurrently, the top 10 ranked variants in the semi-conserved residues of UNG critical motifs (the 4-Pro loop, the uracil recognition β2 strand and the Leu272 loop) involved in enzymatic reactions were also identified. These TDG-nCas9 variants (total 48) were then transfected into HeLa cells together with an sgRNA targeting the Dicer-1 locus, and editing efficiencies were determined through HTS. The process was reiterated by combining the identified variants from the previous round. (**B**) High success rate in validating PLM-predicted TDG variants. The top 38 predicted TDG variants from full-length protein and top 10 variants from the CD domain as shown in (**A**) were inserted into nCas9 protein and transfected into HeLa cells. Five days after the transfection, base substitutions and T>G substitutions were analyzed by HTS. The green circle area highlights the variants showing over two-fold enhancement over the original TDG. (**C**) Identified activity-enhancing variants of TDG exhibited increased activities in three additional loci. Similar to Figure 2B, the top three variants identified from the Dicer-1 locus were further analyzed in the #25, #1, and #14 loci. Nucleotide substitutions were determined using HTS. The presented data are summary of two or three independent experiments, and error bars represent the standard deviation of the mean. (**D**) The top five single residue variants identified in (**B**) were combined and their activities were analyzed at the Dicer-1 site. Data are a summary of two independent experiments. (**E-G**) Summary of TSBE3 EK activities in seven endogenous loci. Frequencies of base substitutions (**E**) and T>S (C and G) conversion purity (**F**) and indel (**G**) at seven endogenous loci were determined using high-throughput sequencing (HTS). The mutated nucleotides were labeled based on their position relative to the 5’ end of the protospacer.

Since even a slight alteration in the evolutionary fitness in a catalytic domain can have profound effects on the function of an enzyme, we independently ranked the variants associated with semi-conservative residues near the critical motifs of the catalytic domain (CD). The top 10 ranked variants were selected for further experimental validation. Despite a relatively low success rate than full-length ranking (**Suppl. Fig.6A**), the examination of these variants identified ones with improved activities for the Dicer-1 site (e.g., R260K), exhibiting a similar magnitude as the L74Q or G107E variants (**Figure 3B**). These improvements were further confirmed on three additional sites (**Figure 3C**).

To further improve the activities of these TDG variants, we combined substitutions that showed the greatest improvements in editing activities from the first rounds, including L74Q, G107E, I37T, H92E, and R260K. Five of the ten combinations outperformed the best single residue variants identified in the first round (**Figure 3D**). The L74Q/G107E and G107E/R260K had the highest activities and performed similarly in most sites we have tested (named as TSBE3 QE and TSBE3 EK, respectively) (**Suppl. Fig. 6B**). Further characterization of TSBE3 QE and TSBE3 EK at additional ten endogenous loci revealed that they predominantly induced T>G and T>C mutations with a purity of T>S (G/C) above 80% (**Figure 3E-F, Suppl. Fig. 6C-D**). Both TSBE3 QE and TSBE3 EK didn’t have a preference over nucleotide sequences surrounding the targeted Ts (**Figure 4A**) and exhibited an editing window spanning protospacer position 3-8 (**Figure 4B**). Notably, despite the increased on-target editing, TSBE3 QE and TSBE3 EK induced comparable low levels of off-target editing at the sgRNA-dependent off-target sites similar to the original TSBE2 (**Figure 4C, Suppl. Table 5**).

**Figure 4.**
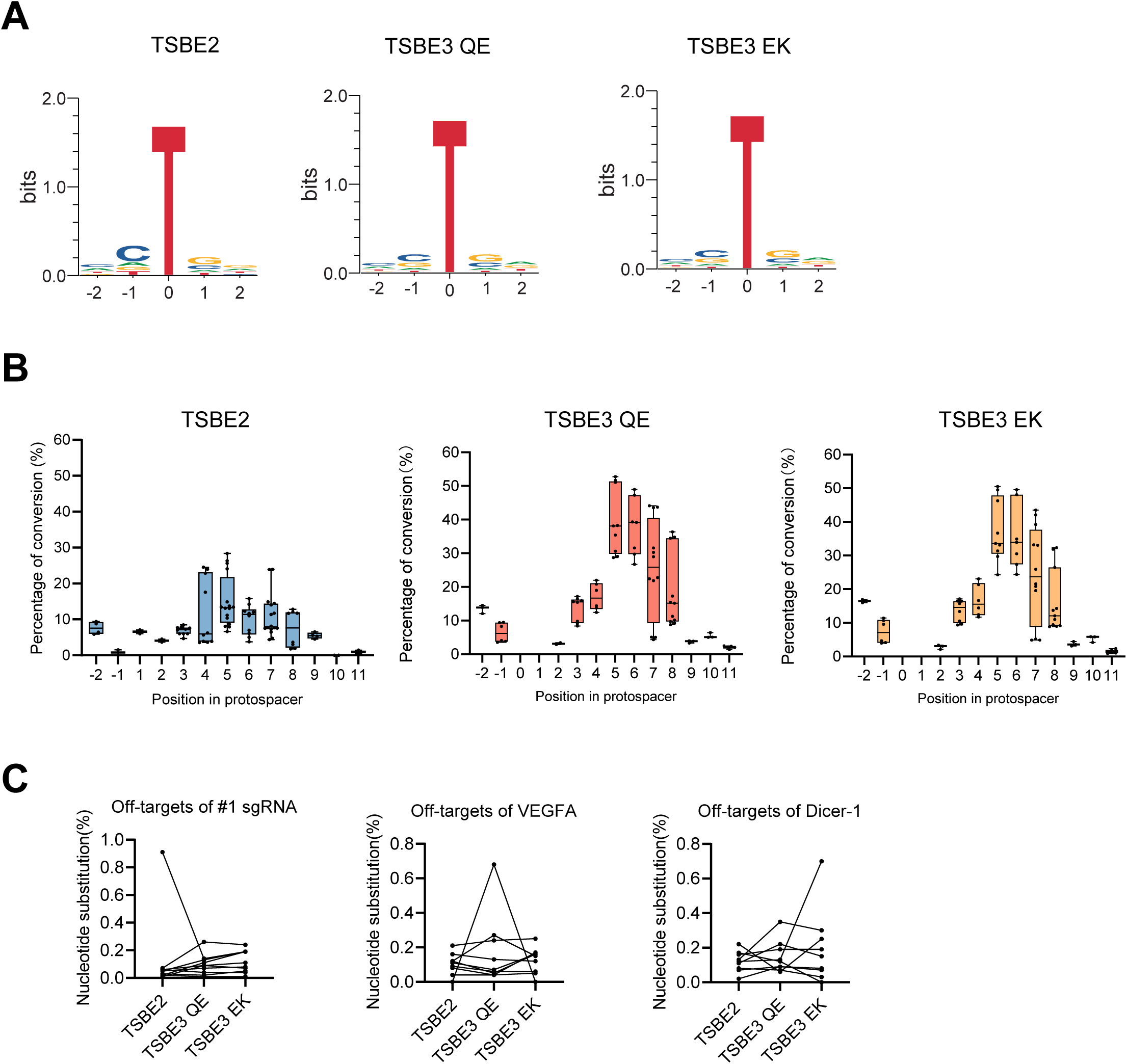
Characterization of TSBE3s. (**A**) Sequence logos representing the composition of nucleotides two base pairs upstream and two base pairs downstream of the edited T bases (position 0) induced by TSBE2 (*left*), TSBE3 QE (*middle*), and TSBE3 EK (*right*). The size of each letter indicates its representation magnitude. (**B**) Editing window for TSBE2 (*left*), TSBE3 QE (*middle*), and TSBE3 EK (*right*). The editing windows were determined by plotting the frequencies of thymine editing across different protospacer positions, with the nucleotide furthest from the PAM designated as position 1. Box plots depict the interquartile range (25th to 75th percentile), horizontal lines represent the median (50th percentile), and whiskers extend to the minimum and maximum values. (**C**) Assessment of off-target activities of TSBEs. HeLa cells were transfected with TSBE2, TSBE3 QE, or TSBE3 EK along with #1 sgRNA (l*eft*), VEGFA (*middle*), or Dicer-1 (*right*) sgRNA. Nucleotide substitutions at predicted off-target sites were determined via high-throughput sequencing (HTS).

### Embedding eTDG between Y1010 and G1011 of nCas9 mitigates indels

Noticeably, the above-characterized glycolysis-based base editors (CGBE-CDG and TSBE3) induced higher frequencies of indels when compared to cytosine or adenine base editors (CBEs or ABEs). This is likely caused by their ability to introduce abasic sites and untimely repair leads to double-stranded breaks. In order to address this issue, we examined the potential of co-expressing HMCES (5-hydroxymethylcytosine binding, embryonic stem cell-specific) with these base editors. HMCES is known to specifically bind to DNA sites containing abasic lesions and form covalent bonds with them, thus safeguarding the DNA from breaks^54^. The co-expression of HMCES with TSBE3 resulted in a significant reduction in the occurrence of indels (**Suppl. Fig. 7A).** Notably, the efficiencies of nucleotide substitution were not affected (**Suppl. Fig. 7B)**. Similarly, the co-expression of HMCES with CGBE-CDG also reduced the generation of indels (**Suppl. Fig. 7C**). These findings suggest that HMCES can effectively mitigate undesirable genetic modifications induced by the glycosylase-based base editors.

Inserting effector modules into different locations of the Cas9 protein can have various effects on the activities of fusion proteins ^41^. We thus investigated whether the insertion of the TDG module at different positions within the nCas9 protein could impact the precision and efficiency of TSBE ^41^. We compared four different inserting locations (F1046 and I1063, Y1010 and G1011, I1029 and G1030, and P1249 and E1250) (**Figure 5A**). We found that inserting TDG EK or TDG TE between nCas9 P1249 and E1250, with a 5-amino acid linker (named TSBE-CE-04-TE), resulted in a shift of the editing window towards the proximal end of the protospacer adjacent to the PAM sequence (**Figure 5B**). For instance, the #14 sgRNAs and TSBE-CE-04-TE predominantly induced base substitutions at T11 instead of T7, and overall editing efficiencies at the targeted loci were not changed. More importantly, this strategy also led to a significant decrease in indels (**Figure 5B**), likely because the abasic sites at different locations of the protospacer may involve distinct repair mechanisms. Similar results were obtained for another two endogenous genomic loci; TSBE-CE-04-TE showed higher efficiencies of nucleotide substitution (**Figure 5C**), higher T-to-G purity of thymine proximal to the PAM sequence (**Figure 5D**), and significantly less indels (**Figure 5E**). Along with our previous results, the co-expression of HMCES with TSBE-CE-04-TE also reduced the generation of indels (**Suppl. Fig. 7D-E**).

**Figure 5.**
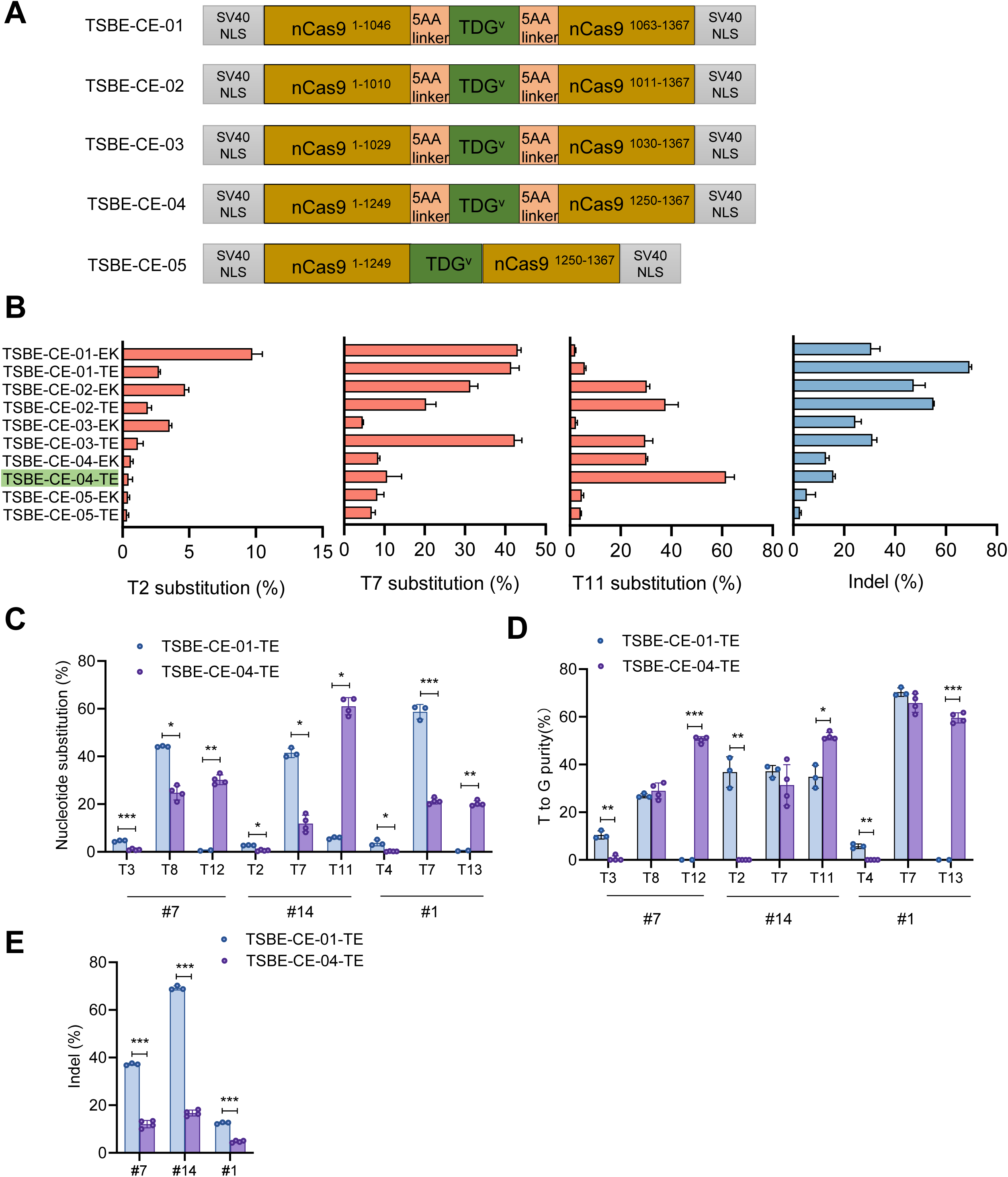
Insertion of TDG into various locations of nCas9 shifts editing window and reduces indels. (**A**). The schematic illustrates the configuration of TDG inserted into various positions in Cas9 protein. TSBE-CE-01, TDG variants inserted between nCas9 F1046 and I1063 with a 5-amino acid linker; TSBE-CE-02, TDG variants inserted between nCas9 Y1010 and G1011 with a 5-amino acid linker; TSBE-CE-03, TDG variants inserted between nCas9 I1029 and G1030 with a 5-amino acid linker; TSBE-CE-04, TDG variants inserted between nCas9 P1249 and E1250 with a 5- amino acid linker; TSBE-CE-05, TDG variants inserted between nCas9 P1249 and E1250 without linker. (**B**). HeLa cells were transfected with TSBEs as shown in (**A**) and the #14 sgRNA. After five days, nucleotide substitution at T2, T7, and T11 as well as indels was determined via HTS. The data shown are summary of two or three independent experiments, and the error bars represent the standard deviation of the mean. (**C-E**). HeLa cells were transfected with TSBE-CE-01-TE and TSBE-CE-04-TE together with indicated sgRNA. Nucleotide substitutions (**C**), T>G purity (**D**), and indels (**E**) were determined by HTS. *, p<0.05, **, p<0.01, ***, p<0.001 in Student’s *t* test.

### TSBE3 precisely corrects db/db mutation in mouse embryos

Having established that TSBE3 could programmably induce T>S substitution, we further tested its capacity to induce nucleotide substitution *in vivo*. We first designed a sgRNA targeting mouse *Dnmt1* T6, mutation of which to C or G leads to synonymous mutation (**Figure 6A**). We delivered the sgRNAs with TSBE3 EK mRNA into mouse embryos by microinjection and transplanted the resulting embryos into surrogate mothers. 12 out of 14 F0 mice (85.7%) harbored base substitutions (**Figure 6B**). Targeted amplicon sequencing of these mice showed that 12 out of 12 mice with base substitutions contained T>C mutation, with frequencies that ranged from 13.4% to 99.5%, and purity ranged from 59.7% to 100% (**Figure 6C-D**). These results indicate that TSBE3 could be utilized to generate murine models with single nucleotide substitutions.

**Figure 6.**
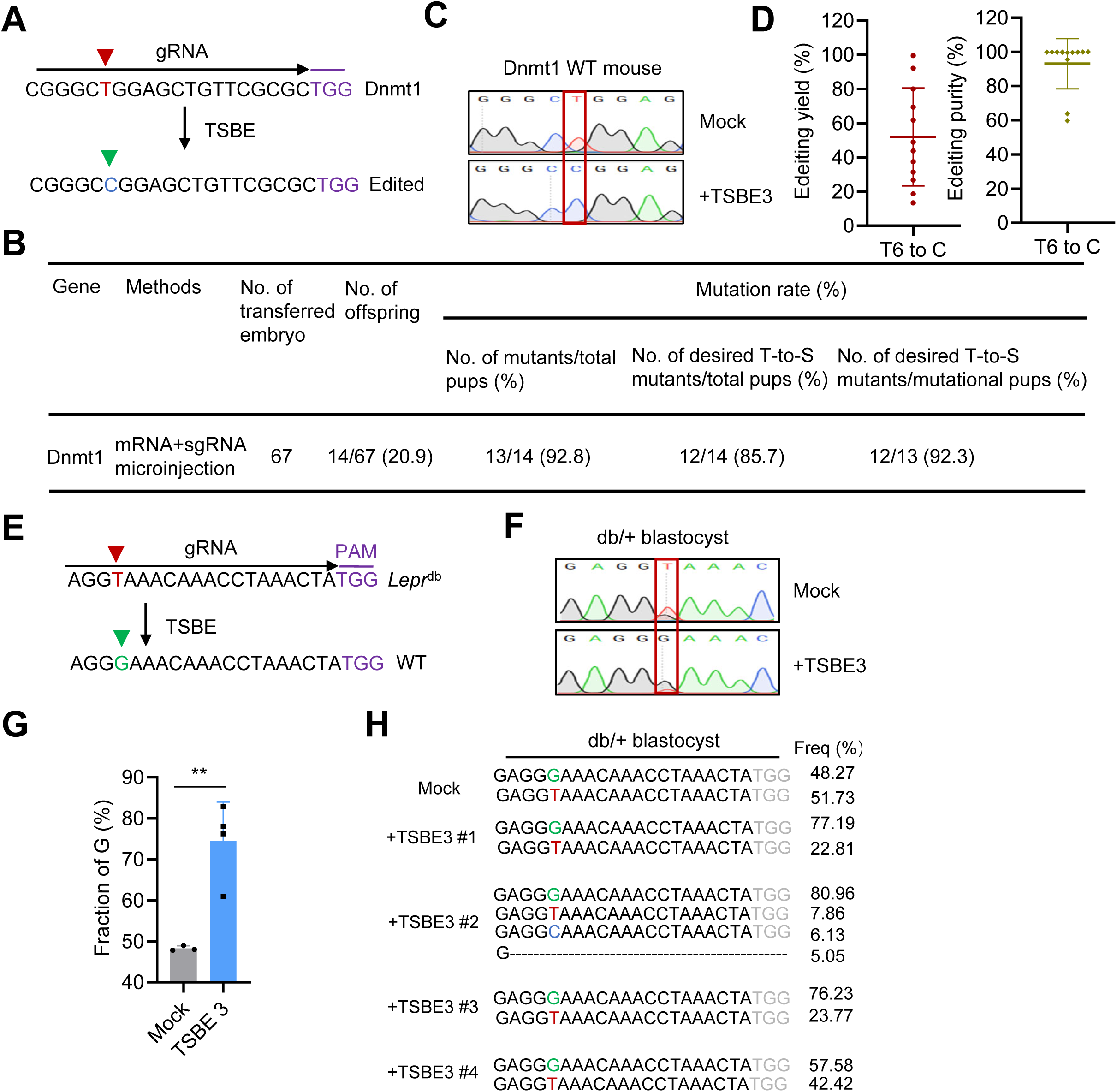
TSBE3 efficiently induces base substitutions in murine embryos. (**A-D**). TSBE3 EK efficiently induced T>S mutation at the *Dnmt1* locus in murine embryos. (**A**) Schematic illustrating the use of TSBE3 EK to induce nucleotide substitution at the *Dnmt1* T6 location. The PAM sequence is displayed in purple, while the targeted thymine in the *Dnmt1* allele is highlighted in red. (**B**). Summary of TSBE3 editing outcomes in 3-week-old pups. WT zygotes of C57B/L6 mice were micro-injected with TSBE3 EK and the sgRNA targeting Dnmt1 T6. A table shows the numbers and frequencies of off-springs, off-springs with intended mutation, and off-springs with T-to-S mutation. (**C-D**). Tail DNA from mice receiving microinjection as in (**B**) were analyzed for nucleotide substitutions and product purity using Sanger sequencing (**C**) and HTS (**D**). Data are representative or summary of the 12 pups with T-to-S mutations. (**E-H**) TSBE3 EK efficiently corrected the *Lepr^db^* mutation in murine embryos. (**E**) Schematic illustrating using TSBE3 EK to correct the *Lepr ^db^* mutation. The PAM sequence is displayed in purple, while the mutated nucleotide in the *Lepr ^db^* allele and the guanine nucleotide corrected by TSBE3 EK are highlighted in red and green, respectively. (**F-G**) db/+ zygotes were obtained *via* in vitro fertilization and were electroporated with a 1:1 quality ratio of TSBE3 EK mRNA (1 ug) and a chemically synthesized sgRNA targeting the *Lepr^db^* mutation (a G>T mutation). Three days after electroporation, blastocysts were collected and analyzed for nucleotide substitutions using Sanger sequencing (**F**) and HTS (**G**). (**H**) Minimal occurrence of editing by-products induced by TSBE3 EK in murine embryos. Mutations induced by TSBE3 EK and their frequencies in db/+ embryos were analyzed using HTS. Data shown are five representative blastocysts.

Furthermore, we employed TSBE3 EK to correct pathogenic G>T mutations in murine embryos. In db/db mice^55^, a common model of obesity, an intronic G-to-T transversion in the leptin receptor gene (*Lepr^db^*) results in mis-splicing and loss of its signal transducing function^56, 57^. To correct this mutation, TSBE3 EK mRNA and a sgRNA targeting the mutation were electroporated into db/+ murine embryos (**Figure 6E**). This resulted in over 55% of the *Lepr^db^* alleles (T allele) being repaired into the WT allele (G allele) (**Figure 6F-G**). Notably, one edited blastocyst had mutations other than the intended T>G substitution, including T>C (∼6%) and indels (∼5%) (**Figure 6H**), while no untended mutations were detected in the other three edited blastocysts (**Figure 6H**). Consistent with the results in the cells lines, no significant off-target editing was observed in the embryos (**Suppl. Table 6**). These findings demonstrate the potential of TSBE to efficiently and precisely reverse pathogenic mutations.

## Discussion

In summary, our study has demonstrated that engineering DNA glycosylases with altered substrate specificities toward normal bases may create orthogonal base editors, which will enable the precise correction of pathogenic mutations and provide essential tools for gene editing and gene therapies. Additionally, the successful engineering of TDG variants highlights the broader utility of protein language models in optimizing biotechnologically relevant enzymes.

Our study demonstrates the potential of protein language models in optimizing enzymes without the need for extensive training data or experimental procedures. By leveraging these models, we were able to obtain variants of UNG with efficient substrate specificities to thymines. It is important to note that only a subset of predicted variants (fewer than 50) was tested in our study, which might have excluded potentially more effective variants. Further investigation of a larger pool of predicted variants could provide a more comprehensive understanding of the enzyme’s capabilities. Additionally, the combination of protein language models and directed protein evolution may further increase their activities as well as their precision.

In contrast to most current base editors, our approach directly fuses the engineered glycosylases (e.g., eTDG) with Cas9 nickase in the absence of deaminases. These findings establish orthogonal strategies for developing novel base editors and highlight the potential of these base editors in gene editing and gene therapies. However, the extent of *in vivo* validation and the evaluation of potential side effects or off-target effects of eTDG (and CGBE-CDG) were not extensively characterized. Future studies should address these aspects to ensure the safety and efficacy of these novel base editors in therapeutic applications.

In conclusion, our study demonstrates the power of protein language models in guiding the engineering of UNG variants for programmable base editing. The enhanced enzymatic activities and programmable capabilities of these novel base editors hold promise for future applications in gene editing and therapy. Further research and optimization of eTDG and similar enzyme variants are warranted to fully explore their potential in precision genome editing.

## Acknowledgments

We thank advanced biomedical technology core facility, supercomputing center, and laboratory animal resource center at Westlake University for the facility support and technical assistance. This work was supported by 2018YFA0801400 (XC) and 2022YFA0807300 (XC) from Ministry of Science and Technology (MOST) of the People’s Republic of China, Key R&D Program of Zhejiang Province (2022SDXHDX0002) (XC); 32025016 & 31870927 (XC) and 81930121 (Y.C.) from National Natural Science Foundation of China (NSFC); 2018YFA0801403 from National Key Research and Development Program of China; 202001BC070001 and 202102AA100053 from Natural Science Foundation of Yunnan Province. This project is supported by Westlake Education Foundation and Tencent Foundation.

## Author contributions

Y.H., C.C, G.C, X.Q.F, and Y.Q.M carried out the experiments. X.B.Z. and F.J.Y. evaluated and optimized the protein language model. Y.H., G.L., and X.C. analyzed the data. Y.H., F.J.Y., and X.C. designed experiments and wrote the paper. All authors read and commented on the paper.

## Material and Method

### Plasmids and vectors

Base editors used in this study were cloned in a pCMV plasmid with blasticidin resistance. sgRNAs were cloned into a pSuper-sgRNA plasmid with puromycin resistance. The wild-type UNG2 sequence (304 aa long) was amplified from cDNA of HEK293T cells, and CDG(UNG2-N204D) and TDG (UNG2-Y147A) were generated via site-directed mutagenesis by PCR, which were then fused with SpCas9 protein as indicated. Protein sequences of CGBEs and TGBEs were listed in Supplementary Table 7. Protospacer sequences of sgRNAs were listed in Supplementary Table 7.

### Cell culture and Transient transfection

HEK293T and HeLa cells were cultured in Dulbecco’s Modified Eagle Medium (DMEM) supplemented with 10% fetal bovine serum (FBS) (Cellmax). Cells were seeded on 48-well plates (Corning) at 2–3×10^5^ cells per well in 500μl of complete growth medium. Between 16 and 24 h after seeding, cells were transfected at 70%–80% confluency with 2.5 μl Polyethylenimine 25000 (Sigma-Aldrich) and 700 ng base editor plasmid, 350 ng sgRNA plasmid. After 24 h of transfection, puromycin (2 μg/ml) and blasticidin (20 μg/ml) were added to the culture for resistance selection. Cells were collected 4 or 6 days after the transfection and were lysed with gDNA lysis buffer (10 mM Tris-HCl, pH 8.0; 0.05% SDS; 25 mg/mL proteinase K (New England BioLabs)) at 37°C for 1 h, followed by enzyme inactivation at 85°C for 30 min.

### Cytotoxicity assay

HeLa cells were seeded in 96-well plates (Corning) at 1×10^4^ cells per well in 250 μl of complete growth medium. 24 hours after seeding, cells were transfected with 1.25 μL Polyethylenimine 25000 (Thermo Fisher Scientific) and 500 ng plasmid containing indicated editors. 0 h, 24 h and 48 h after transfection, 10 μl CCK8 solution was added to each well and the absorbance at 450 nm was measured after 1.5 hours using a microplate reader.

### Target sequencing of endogenous sites in HeLa cells and data analysis

Genomic sites of interest were amplified from genomic DNA and analyzed by HTS as previously described^58^. Briefly, the primary PCR was performed to amplify the targeted genomic DNA with a pair of site-specific primers with common bridging sequences (5’-ggagtgagtacggtgtgc-3’ and 5’-gagttggatgctggatgg-3’) added at the 5’ end. The primary amplification was performed in a 25 µL reaction volume containing 50 ng of gDNA, 0.4 µmol L^−1^ of locus-specific forward and reverse primer, and 12.5 µL 2×Hieff Canace® Plus PCR Master Mix (YEASEN, China). The secondary amplification was conducted in 20 µL preassembled kits, each containing10 µL 2×Taq Master Mix, 200 nmol L^−1^ 2P-F and 2P-R primer, 2 nmol L^−1^ F-(N) and R-(N) primer, and 1 µL primary PCR product. The NGS libraries were sequenced using the Illumina HiSeq platform (Illumina, USA). Sequencing reads were demultiplexed using MiSeq Reporter (Illumina). Alignment of amplicon sequences to a reference sequence was performed using CRISPResso2^59^, a 10-bp window was used to quantify modifications centered around the middle of the 20-bp gRNA. Otherwise, the default parameters were used for analysis. The output files, ‘CRISPResso_quantification_of_editing_frequency.txt’ and ‘Quantification_window_nucleotide_percentage_table.txt’, were combined to calculate the base substitution and indel rates for each individual targeting.

### DNA off-target editing analysis

Cas-OFFinder (<u>CRISPR RGEN Tools (rgenome.net)</u>) used for prediction of potential off-target sites of Cas9 RNA-guided endonucleases, the top 10 off-target sites were selected for validation. Ten potential off-target sites of each target site were amplified from genomic DNA prepared and sequenced on Hi-TOM platform.

### Preparation of sgRNA and mRNA

Chemically modified sgRNAs were synthesized by Genscript, and mRNA of indicated editors was synthesized using an *in vitro* RNA transcription kit (mMESSAGE mMACHINE T7 Ultra Kit, Ambion). In brief, TSBE coding region was cloned in a T7 RNA polymerase promoter plasmid with a 93 poly(A) tail (Supplementary Table 7). T7-TSBE linearized plasmid DNA was transcribed with an in vitro RNA transcription kit (mMESSAGE mMACHINE T7 Ultra Kit, Ambion) following the manufacturer’s instructions. TSBE mRNA was eluted in nuclease-free water and stored at −80 °C.

### Animal care, microinjection and electroporation of zygotes

All animal experiments were in compliance with regulations by the Association for Assessment and Accreditation of Laboratory Animal Care in Hangzhou and were ratified by the Laboratory Animal Resources Center of Westlake University. Db/db mice were obtained from the Shanghai Model Organisms, and WT C57BL/6J and CBA/J mice were obtained from the Laboratory Animal Resources Center of Westlake University. Mice were housed in a specific pathogen-free facility on a 12-h light and 12-h dark cycle with ample access to food and water. For microinjection, WT C57BL/6J mouse strains were used as sperm and egg donors. The mixture of TGBE mRNA (100 ng μl^−1^)/sgRNA (100 ng μl^−1^) was diluted in Opti-MEM media (Gibco) and injected into cytoplasm using an Eppendorf TransferMan NK2 micromanipulator. Injected zygotes were transferred into pseudopregnant female mice immediately after injection or after overnight culture in KSOM medium at 37 °C under 5% CO2 in air. For electroporation, db/db and WT C57BL/6J mouse strains were used as sperm and egg donors respectively. Zygotes were pipetted into 10 μl drops of Opti-MEM media (Gibco) and the entire content mixed with 10 μl of the TGBE mRNA (100 ng μl^−1^)/sgRNA (100 ng μl^−1^) reconstituted in TE buffer (pH 7.5) and deposited into a 1 mm electroporation cuvette (Harvard Apparatus) and electroporated using a ECM830 Square Wave Electroporation System (BTX Harvard Apparatus) with 30 V, 1 ms pulse duration and two pulses separated by 100 ms pulse interval. Following the electroporation, a pre-warmed 100 μl aliquot of KSOM/BSA media was added into the cuvette and zygotes were removed from the cuvette with a sterile plastic pipette, which were then cultured to blastocysts for three days (37°C and 5% CO_2_).

### Statistical analysis

Statistical tests performed by GraphPad Prism 8 included the two-tailed, unpaired, two-sample t-test or Dunnett’s multiple comparisons test after One-Way ANOVA. Data are presented as mean ± s.d. from independent experiments.

### Developing PLM-based strategies for enhancing enzymatic activities

Seventeen PLMs and nine ranking strategies were evaluated to determine the best PLM-based strategies for enzymatic evolution. Briefly, we first define the wildtype sequence and mutated sequence at the i-th site as 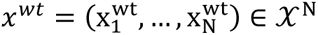 and 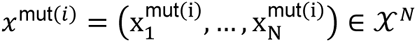 respectively. 𝒳 represents the set of amino acids, N denotes the length of the sequence and 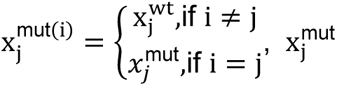 refers to the mutation at j-th site.

A collection of masked language models, denoted as p(x|z), were employed to enable the computation of the probability of a target sequence x (representing either a wildtype or mutated sequence) given a background sequence z (typically with certain masked regions). Various rank strategies are developed based on distinct choices of background sequences for the wildtype sequence x^wt^ and mutated sequence at i-th site, denoted as x^mut(i)^.

All rank strategies employ a nearly identical score function that evaluates the difference in logarithmic probabilities at each site *k* across the sequence length *N*. Specifically, this score function compares the mutated sequence at the i-th site, denoted as 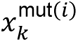, given a mutated background 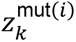, with the wildtype sequence 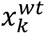 given a wildtype background 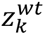.

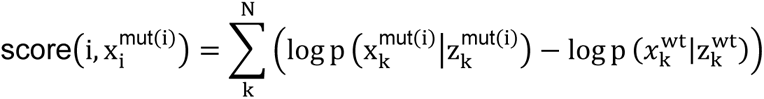

A prior investigation^53^ introduced three distinct backgrounds, namely Masked Marginal Probability (esm1v_1, esm1v_2, esm1v_3), Mutant Marginal Probability (esm1v_4), and Wildtype Marginal Probability (esm1v_5). The Masked Marginal Probability (esm1v_1, esm1v_2, esm1v_3) approach utilizes a masked background sequence for calculating the probabilities of target sequences. Based on different masking strategies, we introduce three specific strategies: Strategy A (esm1v_1), Strategy B (esm1v_2), and Strategy C (esm1v_3). Detailed explanations of these strategies will be provided in the subsequent sections.

The Mutant Marginal Probability (esm1v_4) approach employs a mutant sequence as the background sequence for calculating the probabilities of target sequences. On the other hand, the Wildtype Marginal Probability (esm1v_5) approach utilizes a wildtype sequence as the background sequence for calculating the probabilities of target sequences.

In addition to the aforementioned approaches, we also introduce another set of strategies referred to as Auto Regressive (AR). Autoregressive techniques have found widespread adoption in the field of Natural Language Processing^60^, where the generation of new tokens is conditioned on previously generated tokens. Inspired by this concept, we have developed four Autoregressive strategies: Strategy A (AR_1), Strategy B (AR_2), Strategy C (AR_3), and Strategy D (AR_4).

As commonly known, the set of amino acids is denoted by 𝒳, where |𝒳| = 20. In our extended framework, we introduce a mask token, denoted as x^mask^ and include it in the set of amino acids, resulting in 𝒳_+_, which comprises the 20 amino acids along with the mask token. Therefore, |𝒳_+_| = 21.

The following nine scoring methods were utilized in the study.

1. MaskMP_A In Masked Marginal Probability Strategy A (MaskMP_A), the background sequence for both the wildtype and mutated sequences consists of full mask tokens throughout the sequence.

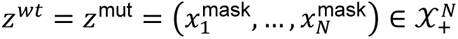 The score function:

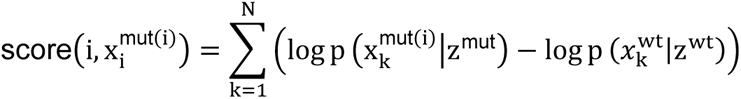
2. MaskMP_B In Masked Marginal Probability Strategy B (MaskMP_B) and Masked Marginal Probability Strategy C (MaskMP_C), a similar mask strategy is employed, where the mutated site i in the mutated sequence, denoted as x^mut(i)^, is masked. However, there is a distinction in the choice of background sequences between the two strategies. MaskMP_B utilizes the target sequences themselves as the background, while MaskMP_C employs the mutated sequence as the background for both the mutated and wildtype sequences.

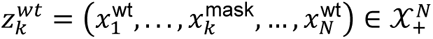

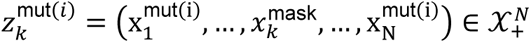 The score function:

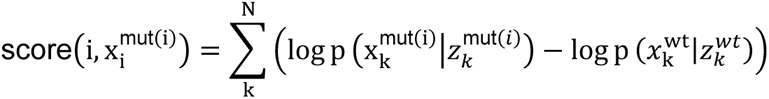
3. MaskMP_C As previously discussed, in Masked Marginal Probability Strategy C (MaskMP_C), the same background sequence is employed for both the mutated and wildtype sequences.

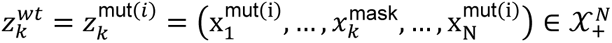 The score function:

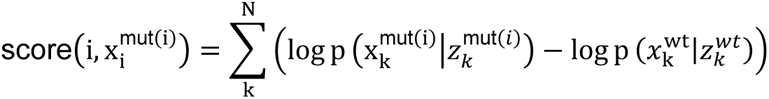
4. MutantMP In Mutant Marginal Probability (MutantMP), the mutated sequence is utilized as the background for both the mutated and wildtype sequences.

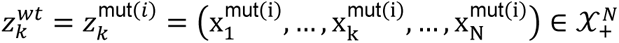 The score function:

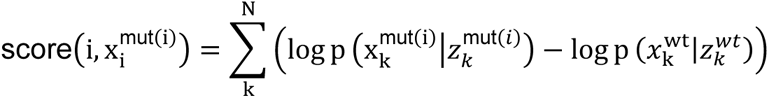
5. WildtypeMP In Wildtype Marginal Probability (WildtypeMP), the wildtype sequence serves as the background for both the mutated and wildtype sequences.

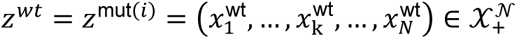 The score function:

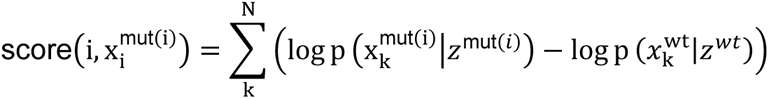
6. AR_A In Autoregressive Strategy A (AR_A), the wildtype sequence is used as the sequence to be replaced, and the mutated sequence is used to replace the wildtype sequence. Notably, both backgrounds in this strategy are the same.

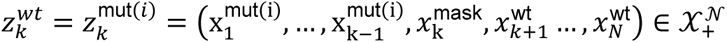 The score function:

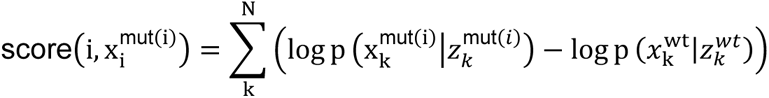
7. AR_B In Autoregressive Strategy B (AR_B), the wildtype sequence is used both as the sequence to be replaced and as the background sequence for the wildtype sequence replacement. Additionally, the mutated sequence is used as the background sequence for the replacement of the wildtype sequence.

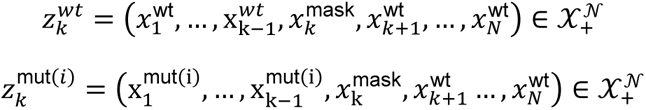 The score function:

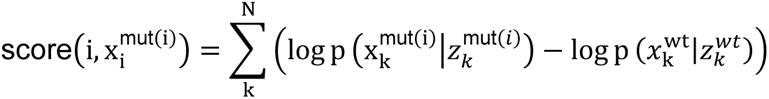
8. AR_C In Autoregressive Strategy C (AR_C), a fully masked sequence is used as the sequence to be replaced, and the mutated sequence is used to replace it. Notably, both backgrounds in this strategy are the same.

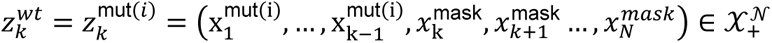 The score function:

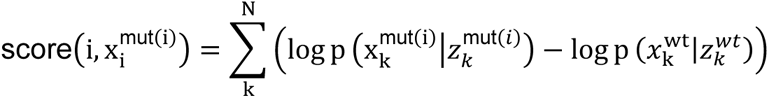
9. AR_D In Autoregressive Strategy D (AR_D), a fully masked sequence is used as the sequence to be replaced. The wildtype sequence is used as the background sequence for replacing the wildtype sequence, while the mutated sequence serves as the background sequence for replacing the masked sequence.

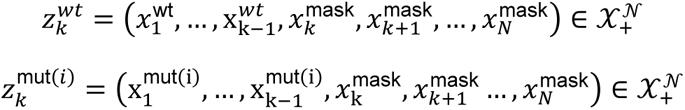 The score function:

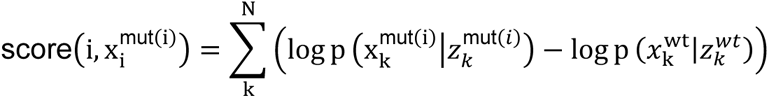

## Supplementary Figures

**Suppl. Figure S1.**
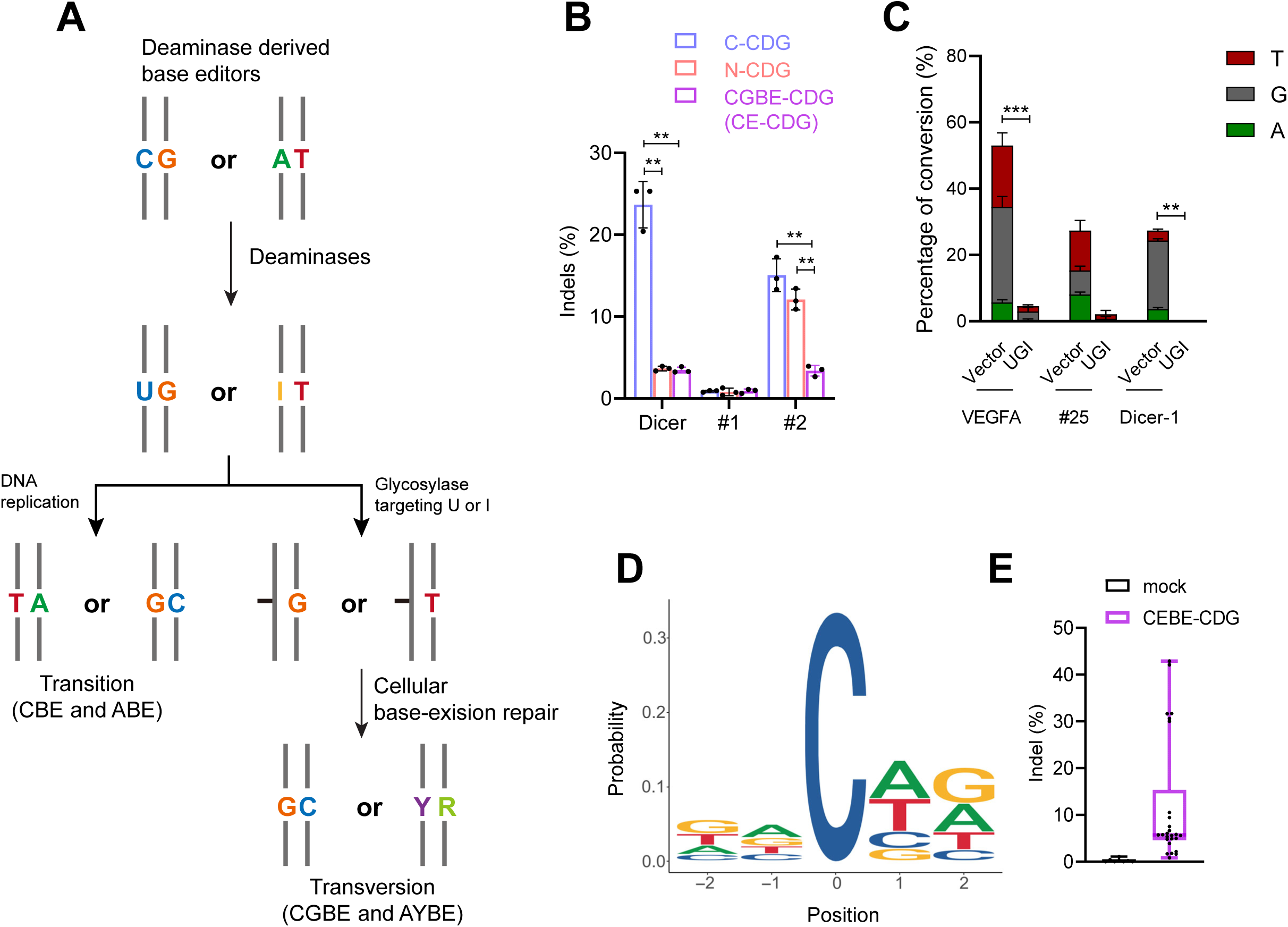
Characterization of CGBE-CDG. (**A**) A diagram showing the principles of current base editors. Current base editors utilize cytidine deaminases (CBE) or adenine deaminases (ABE) to programmably generate modified bases (uracil or hypoxanthine respectively) in targeted loci, which upon replication result in transition mutations (C>T or A>G). Alternatively, abasic sites can be induced by including specific glycosylases against the modified bases (e.g., UNG or MPG), leading to transversion mutations. (**B**) CE-CDG induces less indels than either N-CDG or C-CDG. As in Figure 1E, HeLa cells were transfected with nCas9-CDG fusion proteins with indicated configurations and Dicer-1, #1, or #2 sgRNAs. The indels (insertions + deletions) generated within a 40 bp range upstream and downstream of the protospacer were determined with HTS. CE-CDG, Cas9-embedded CDG; N-CDG, CDG was fused to the N-terminus of nCas9 protein; C-CDG, CDG was fused to the C-terminus of nCas9 protein. (**C**) CE-CDG requires the enzymatic activity of the UNG variant. HEK293T cells were transfected with CE-CDG and Dicer-1 or VEGFA or #25 sgRNA with or without UGI. Seven days after the transfection, the editing efficiencies were determined with HTS. Data are summary of two or three independent experiments. (**D**) Composition of nucleotides two bps upstream and two bps downstream of the mutated C bases (position 0) were calculated and plotted as sequence logo. The size of each letter indicates the magnitude of representation. (**E**) Summary of indels generated by CE-CDG in seven endogenous loci. As in Figure 1F, the indels (insertions + deletions) generated within a 40 bp range upstream and downstream of the protospacer were determined with HTS. (**B, C, D, E**) Error bars stand for the standard deviation of the mean. **, p<0.01, ***, p<0.001 in Student’s *t* test.

**Suppl. Figure S2.**
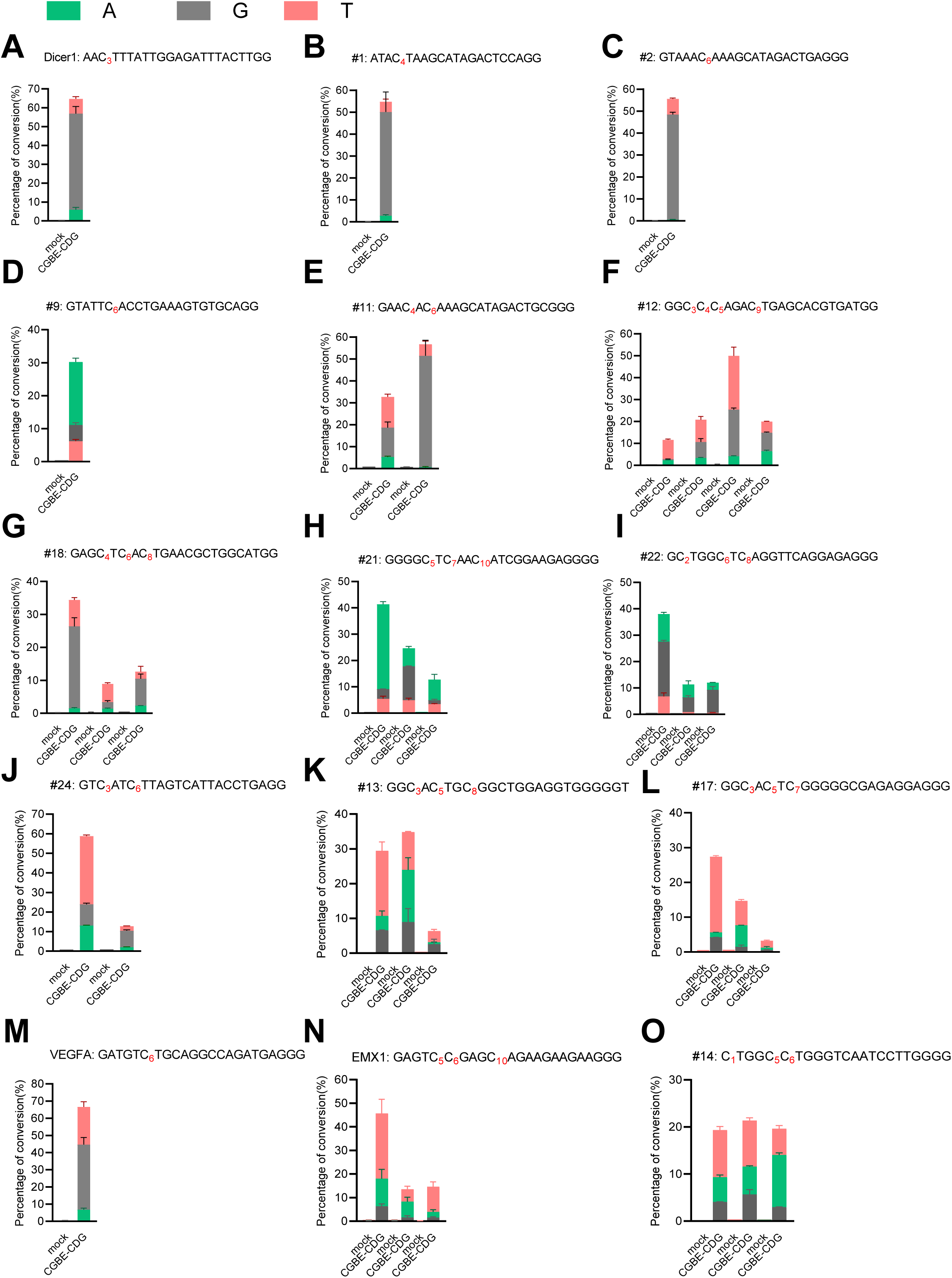
Activities of CGBE-CDG in 15 endogenous loci. (**A-O**) HeLa cells were transfected with the indicated sgRNA and CGBE-CDG at a 1:1 molar ratio. Seven days after the transfection, nucleotide substitution was determined with HTS. Nucleotides with detectable mutation (>0.2%) were highlighted and numbered starting from the position most distal to the PAM. Data are summary of two-four independent experiments. Error bars stand for the standard deviation of the mean.

**Suppl. Figure S3.**
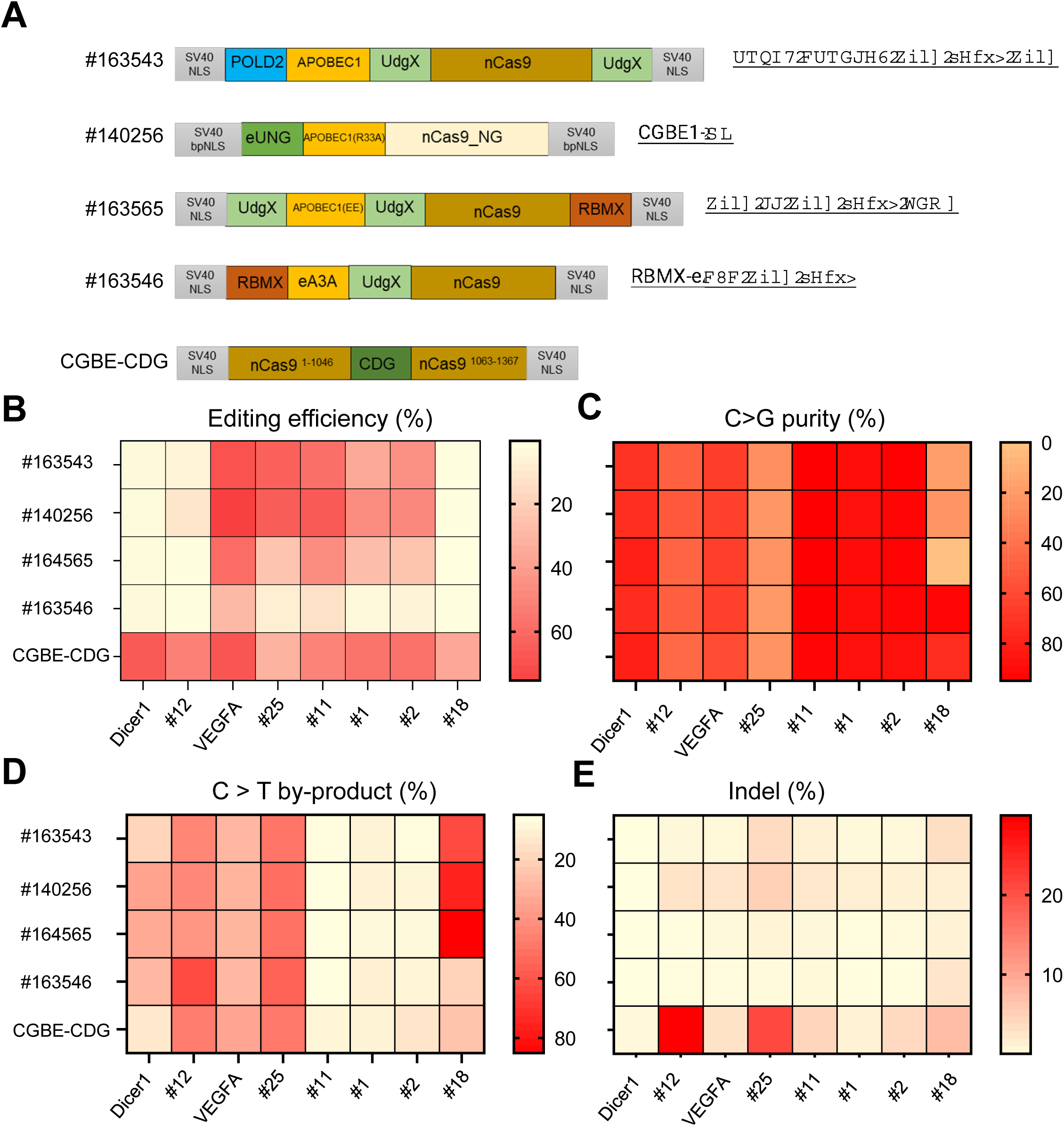
Comparison between CGBE-CDG and previously reported CGBEs. (**A**) Schematics of previously reported CGBE^23–25^, including POLD2-APOBEC1-UdgX-nCas9-UdgX (addgene #163543), CGBE1-NG (Addgene #140256), UdgX-EE-UdgX-nCas9-RBMX (addgene #163565), and RBMX-eA3A-UdgX-nCas9 (Addgene #163546). (**B-E**) CGBEs as indicated in (**A**) were co-transfected along with indicated sgRNAs into HeLa cells. Seven days following the transfection, the target loci were amplified and analyzed with HTS. Each square represents the average of three independent experiments. (**B**) Heat map illustrating the percentages of nucleotide substitutions (C to T% + C to G% + C to A%) at eight endogenous loci. (**C**) As in (**B**), heat map showing C>G purity of the editing products (C to G/(C to T + C to G + C to A)%) at eight endogenous loci. (**D**) Heatmap showing the ratios of the C-to-T byproduct (C to T/(C to T + C to G + C to A)%) at eight genetic loci. (**E**) as in (**B**), a heatmap showing the indels (insertions + deletions) produced within a 40 bp range upstream and downstream of the editing window at the eight genetic sites.

**Suppl. Figure S4.**
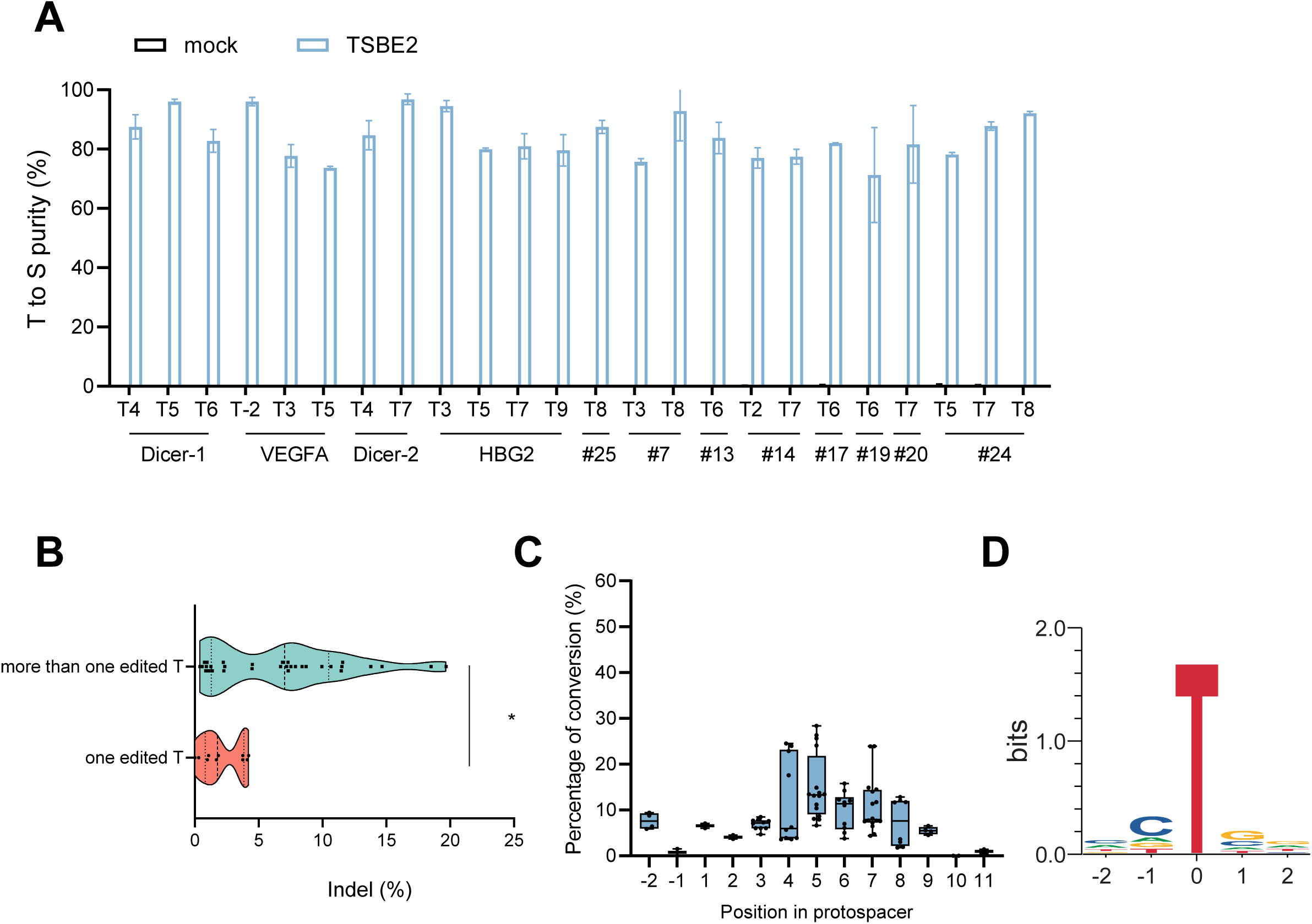
Characterization of TSBE2. (**A**) Summary of T>S purity of TSBE2 in 12 endogenous loci. HeLa cells were transfected with the indicated sgRNA and TSBE2 at a 1:1 molar ratio. Seven days after the transfection, nucleotide substitution was determined with HTS. (**B**) Comparison of indels produced by TSBE2 in 12 endogenous loci, 12 endogenous loci are divided into two categories, one with only one edited T within the editing range, and the other with multiple edited Ts within the editing range. (**C**) The editing window for TSBE2 was determined by plotting the frequencies of cytosine editing across various protospacer positions, with the nucleotide furthest from the PAM designated as position 1. Box plots represent the interquartile range (25th to 75th percentile), horizontal lines indicate the median (50th percentile), and whiskers extend to the minimum and maximum values. (**D**) Composition of nucleotides two bps upstream and two bps downstream of the mutated T bases (position 0) were calculated and plotted as sequence logo. The size of each letter indicates the magnitude of representation.

**Suppl. Figure S5.**
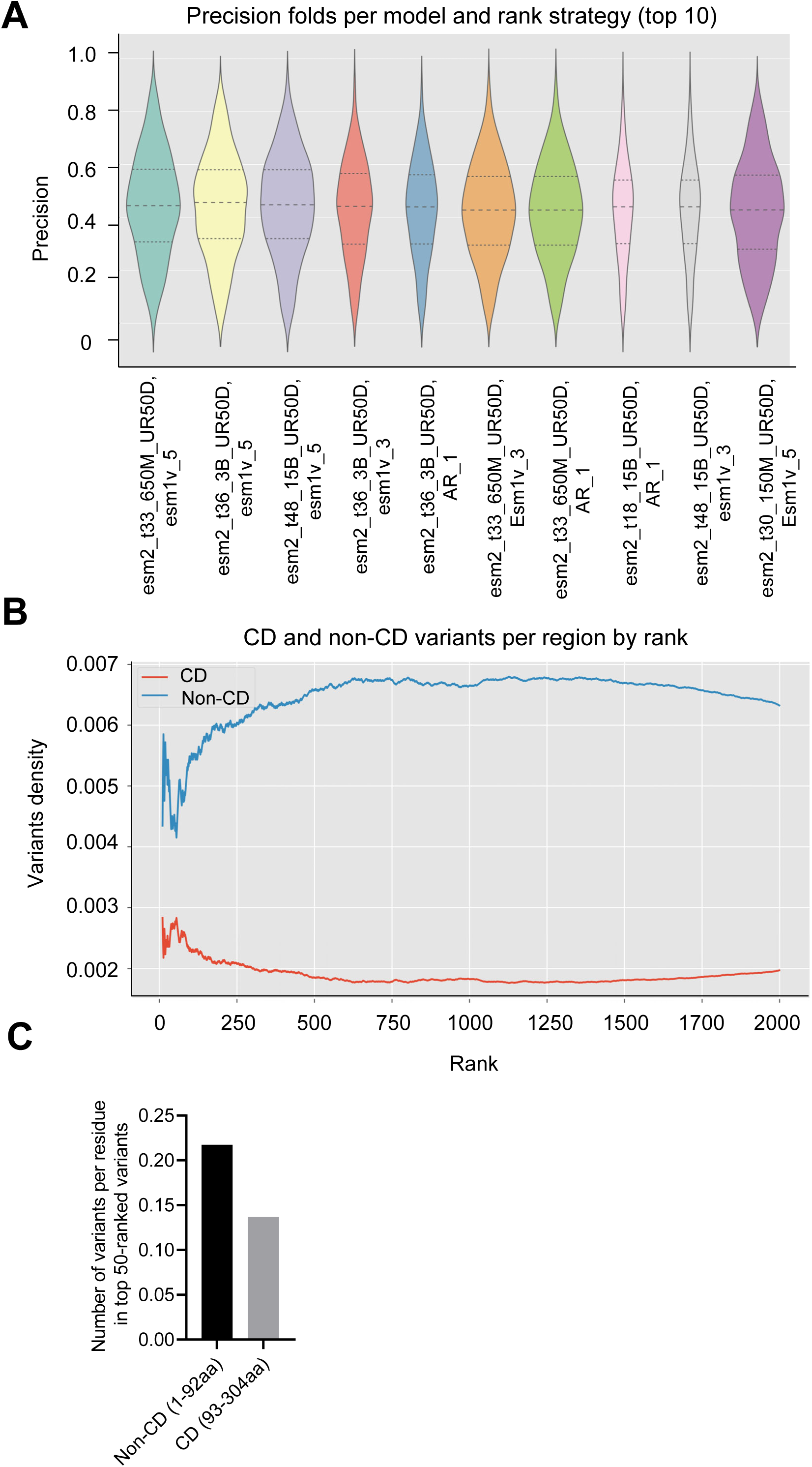
Developing protein language models and ranking strategies for predicting function-augmenting variants. (**A**) Seventeen language models and nine ranking strategies were assessed using a previously published dataset to identify the most effective strategies for predicting function-enhancing variants. To generate the predicted positive samples, the top 50 variants for each scoring method were selected, as well as any variants that scored higher than the wildtype. Real positive samples consisted of variants with a fitness level greater than that of the wildtype. The precision of the prediction was calculated as the proportion of real positive samples among the predicted positive samples. The precision of the top ten strategies was graphed. (**B**) PLM-ranked function-enhancing variants were enriched in the N-terminal tail of TDG. The density of TDG variants with substitutions in the CD region and non-CD region was calculated ranking from 10 to 2000. (**C**) Enrichment of the top 50 ranked TDG variants in the N-terminus. The number of variants per residue in the top 50 ranked variants was calculated in either the non-CD (N-terminus, 1-92aa) or the CD domain (93-304 aa).

**Suppl. Figure S6.**
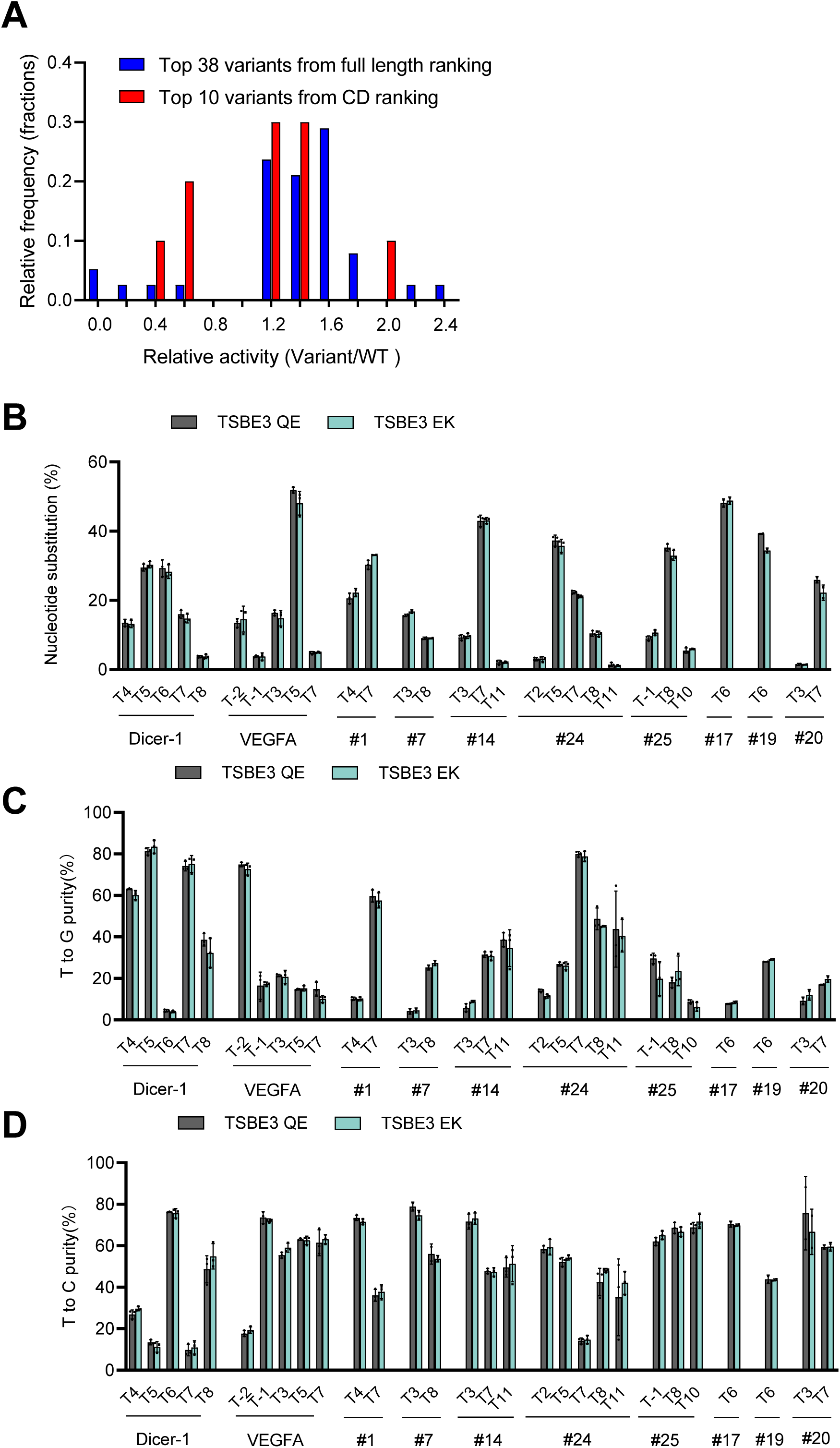
Obtaining highly efficient TDG variants assisted with PLM. (**A**) PLM enables the identification of augmented TDG variants with a high success rate. The histogram shows the distribution of activities among the analyzed TDG variants relative to the wild type. To assess the activities of the mutant TDGs, they were fused with nCas9 and transfected into HeLa cells together with Dicer-1 sgRNA. Nucleotide substitutions of T5 were determined by HTS. The y-axis represents the frequency of TDG variants, while the x-axis represents the range of enzyme activity levels. (**B-D**) Comparison of TSBE3 QE and TSBE3 EK in ten endogenous loci. HeLa cells were transfected with TSBE3 QE or TSBE3 EK along with the indicated sgRNAs. Nucleotide substitutions (**B**), T>G purity (**C**), and T>C purity (**D**) were determined via HTS. Error bars represent the standard deviation of the mean, and the data presented are summary of two or three independent experiments.

**Suppl. Figure S7.**
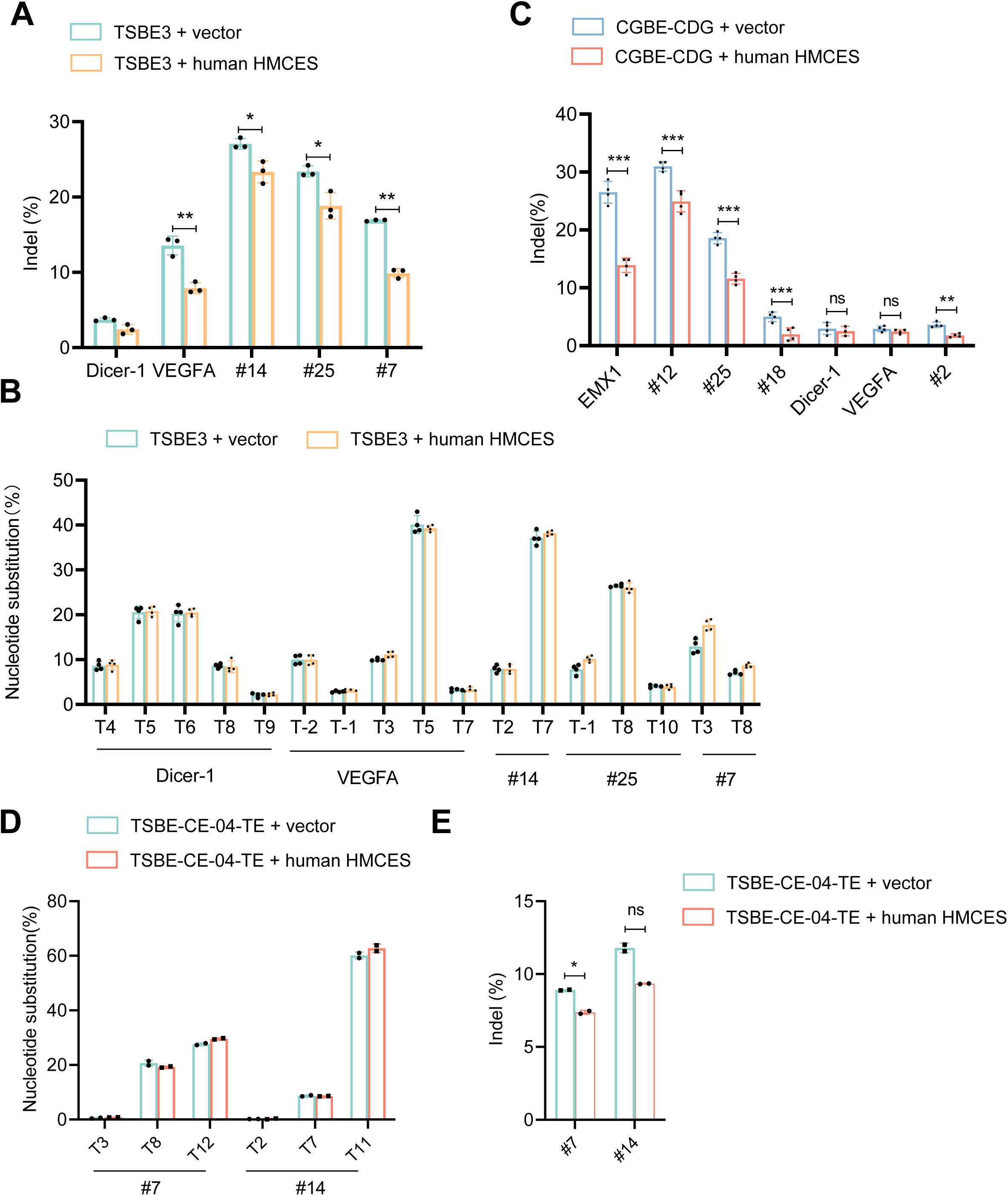
HMCES mitigates indels induced by CGBE-CDG and TSBE3. (**A**) HeLa cells were transfected with TSBE3 and corresponding sgRNAs, either with or without a vector expressing human HMCES. Indels were assessed through high-throughput sequencing (HTS) five days after the transfection. (**B**) As in (**A**), frequencies of nucleotide substitution were determined at five endogenous sites in HeLa cells following the transfection of TSBE3 and corresponding sgRNAs, with or without co-expressing HMCES. (**C**) HeLa cells were transfected with CGBE-CDG and corresponding sgRNAs, either with or without a vector expressing human HMCES. Indels were assessed through high-throughput sequencing (HTS) five days after the transfection. (**D-E**) HeLa cells were transfected with TSBE3 and corresponding sgRNAs, either with or without a vector expressing human HMCES. Frequencies of nucleotide substitution (**D**), indels (**E**) were determined through high-throughput sequencing (HTS) five days after the transfection. (**A-E**) Data are summary of two or four independent experiments, and error bars represent the standard deviation of the mean. *, p<0.05, **, p<0.01, ***, p<0.001 in Student’s *t* test.

**Suppl. Table 1. Summary of CGBE-CDG-induced substitution and indels in predicted off-target sites.** Cytidines with detected substitutions (>0.1%) are highlighted in red, and PAM sequences are shaded in gray. The data presented represent the average of two independent experiments. Importantly, none of the off-target sites exhibited editing activities above 0.2%.

**Suppl. Table 2. A list of protein language models and ranking strategies evaluated in this study.**

**Suppl. Table 3. Performance of different PLM and ranking strategies on a previously published dataset.**

**Suppl. Table 4. Predicted fitness of the top 1000 TDG single residue variants.**

**Suppl. Table 5. Summary of TSBE3 EK and TSBE3 QE-induced substitutions and indels in predicted off-target sites.** Thymines with detected substitutions (>0.1%) are shown in red, and PAM sequences are shaded in gray. The data represented the average of two or three independent experiments. Notably, none of the off-target sites exhibited editing activities above 1%.

**Suppl. Table 6. Summary of TSBE3-induced mutations in predicted off-target sites in murine embryos.** Mutation frequencies were determined for all Ts in the top ten off-target sites, as depicted in Figure 2f. PAM sequences are shaded in gray. The data represent the average of four db/+ blastocysts that were corrected using TSBE3.

**Suppl. Table 7. Sequences of sgRNAs and base editors utilized in this study.**

